# ChronoRoot: High-throughput phenotyping by deep segmentation networks reveals novel temporal parameters of plant root system architecture

**DOI:** 10.1101/2020.10.27.350553

**Authors:** Nicolás Gaggion, Federico Ariel, Vladimir Daric, Éric Lambert, Simon Legendre, Thomas Roulé, Alejandra Camoirano, Diego H. Milone, Martin Crespi, Thomas Blein, Enzo Ferrante

## Abstract

**Background:** Deep learning methods have outperformed previous techniques in most computer vision tasks, including image-based plant phenotyping. However, massive data collection of root traits and the development of associated artificial intelligence approaches have been hampered by the inaccessibility of the rhizosphere. Here we present ChronoRoot, a system which combines 3D printed open-hardware with deep segmentation networks for high temporal resolution phenotyping of plant roots in agarized medium.

**Results:** We developed a novel deep learning based root extraction method which leverages the latest advances in convolutional neural networks for image segmentation, and incorporates temporal consistency into the root system architecture reconstruction process. Automatic extraction of phenotypic parameters from sequences of images allowed a comprehensive characterization of the root system growth dynamics. Furthermore, novel time-associated parameters emerged from the analysis of spectral features derived from temporal signals.

**Conclusions:** Altogether, our work shows that the combination of machine intelligence methods and a 3D-printed device expands the possibilities of root high-throughput phenotyping for genetics and natural variation studies as well as the screening of clock-related mutants, revealing novel root traits.

## Background

Plants are sessile organisms unable to seek out optimal environmental conditions for development and survival. Strikingly, a remarkable developmental plasticity allows plants to complete their life cycle under changing growth conditions [1]. Understanding plant root plastic growth is crucial to assess how different populations may respond to the same soil properties or environmental conditions and to link this developmental adaptation to their genetic background [2]. Under controlled conditions, root development is generally observed based on images of plants growing vertically on the surface of a semisolid agarized medium. Root system architecture (RSA) is then characterized by parametrization of a grown plant, which relies on the combination of a subset of variables like main root (MR) length or density and length of the lateral roots (LRs) [3]. Several semi-automatic tools have been developed to assist root phenotyping at specific time points [4]. However, temporal phenotyping is generally hindered by technological limitations, ignoring potentially useful phenotypical parameters that may be linked to the temporal dynamics of root growth. Here we present ChronoRoot, a low-cost system based on off-the-shelf electronics, 3D printed hardware components and deep learning models, allowing high-throughput temporal phenotyping of *Arabidopsis thaliana* RSA. Figure 1 illustrates the different components of ChronoRoot. Temporal sequences of pictures, automatically snapped, are processed for root segmentation through a convolutional neural network (CNN) model. We leverage the latest advances in CNNs for image segmentation and propose an architecture for RSA delineation which incorporates deep supervision, producing fast and accurate segmentations. The root extraction workflow is completed by a temporal consistency refinement step and a final graph generation process, which generates a labeled root graph for every image. An exploratory approach assessing root growth under alternative photoperiods served to demonstrate that temporal phenotyping performed by ChronoRoot allows deciphering the evolution of the traditional RSA parameters throughout time. Moreover, novel parameters emerged, including architectural features, oscillating growth speed and other characteristics derived from spectral analysis of the growth signals in the Fourier domain. The combination between a low-cost automatic device for image acquisition and machine intelligence methods for image segmentation gave rise to a powerful tool for root phenomics potentially applicable to natural variation studies, the characterization of root-related subtle disorders and the screening for clock-associated mutants.

**Figure 1.**
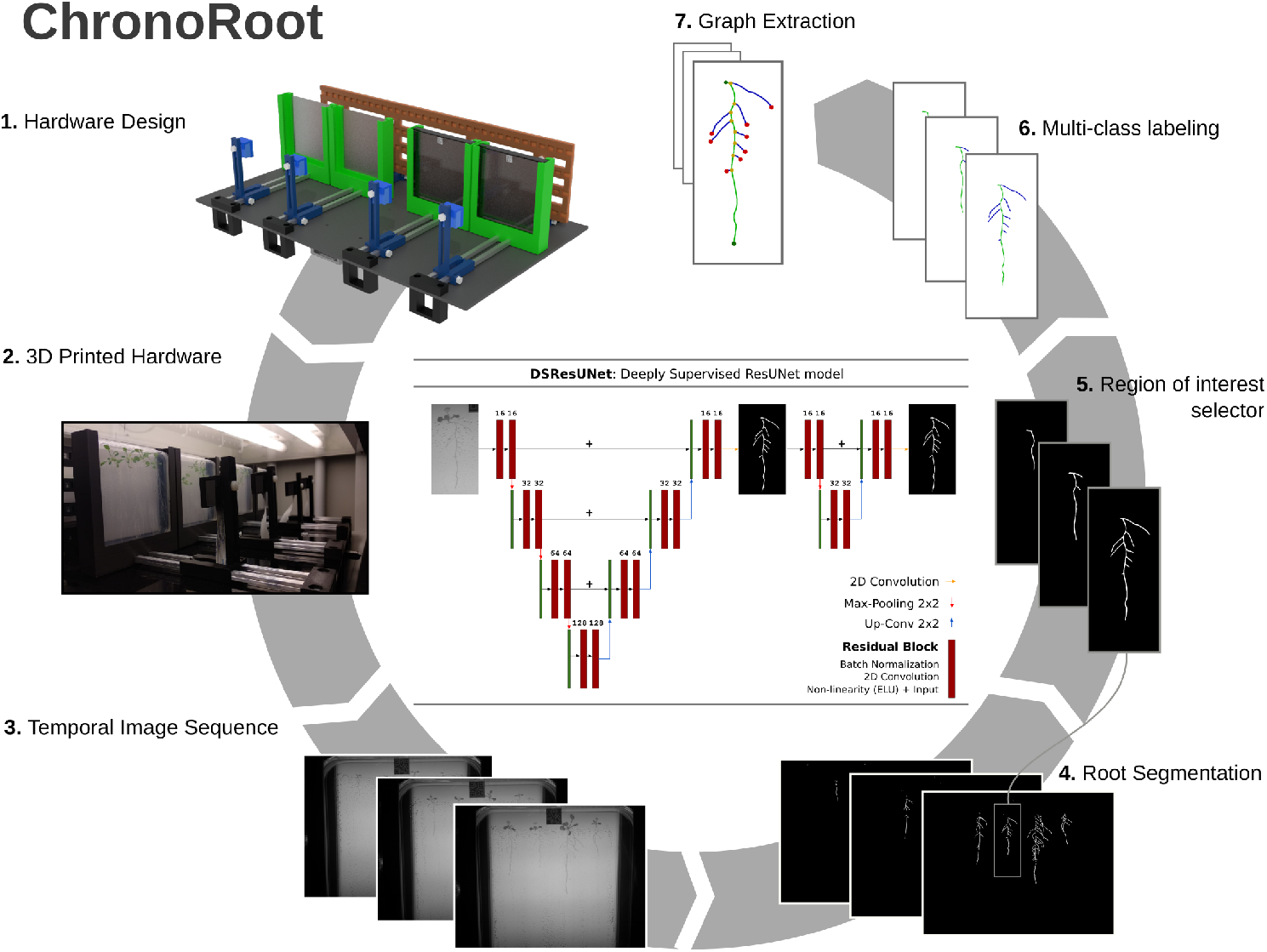
Main components of ChronoRoot. 1) Open hardware specification (see the Supplementary File 1 for a detailed description). 2) 3D printed ChronoRoot system mounted in a plant growth chamber. 3) Temporal sequence of images acquired by the system are provided as input to the CNN based segmentation module. The diagram corresponding to the proposed CNN architecture is included in the center of the Figure. 4) The deep learning model produces dense segmentation maps for all the plants, which are enhanced taking into account the temporal consistency of the results. 5) Independent plants can be selected to be processed individually. 6,7) The roots are skeletonized and a graph is constructed by traversing the skeleton. Pixels in the skeleton are identified as belonging to the main root (green) or lateral root (blue). The graph nodes are labeled as being the main root seed and tip (green), lateral root tip (red) or bifurcation (yellow).

## Data Description

### Plant material and growth conditions

*Arabidopsis thaliana* ecotype Col-0 seeds were surface sterilized and stratified at 4°C for 2d before being grown under long day conditions (16h light, 140μ*Em*^−2^*s*^−1^/ 8h dark), or continuous light (24h light, 140μ*Em*^−2^*s*^−1^) at 22°C, on half-strength Murashige and Skoog media (1/2 MS) (Duchefa, Netherlands) with 0.8% plant agar (Duchefa, Netherlands). Four seeds were used per plate. All the experiments where performed under laboratory conditions according to the local institutional guidelines.

### Datasets

We generated two different datasets in this work: the first one was used to train and evaluate the segmentation performance of the CNN models, while the second one served as an exploratory use case, to assess root growth under alternative photoperiods and provide an example of the novel temporal phenotypical parameters that can be extracted with ChronoRoot. Note that all these images were obtained with the proposed 3D printed hardware, and both are available to encourage reproducible research. Importantly, when splitting the training, validation and test partitions, we were careful not to include images corresponding to the same video on different partitions, to avoid overoptimistic biased evaluations.

- **Dataset used to train and validate the deep learning models for root segmentation:** The dataset used for training consisted of 331 images from 55 videos (on average 6 images from the same plate at different states of growth), 11 of those videos were annotated by an expert biologist. The dataset used for testing consisted of 55 images from 11 different videos, all annotated by the same expert. The tool used for the manual annotation was ITK-SNAP[5]. In total, 240 plants distributed over the 66 videos were used for training/testing the methods.
- **Use case dataset for plant phenotyping under alternative photoperiods:** We used 12 videos for each photoperiod, with pictures taken every 15 minutes. We took the first 17 days (1632 frames), and after processing the videos we proceeded to discard the results from the first 3 days prior to seed germination. We selected 25 plants from each photoperiod to perform the temporal growth analyses.

## Analyses

We designed an automatic method to perform RSA delineation in temporal image sequences of plant roots. Our framework takes a sequence of images as input and outputs a labeled graph for each frame, representing the current root growing state. Graphs are powerful data structures particularly useful to represent curvilinear shapes like plant roots (details on the graph generation process are provided in the Methods section). The main module of the RSA delineation method is a deep CNN which produces a dense segmentation mask, where every pixel is classified as belonging to the root or the background. We proposed different CNN architectures for this task (described in the Methods section), and compared their performance with state-of-the-art models using manual annotations produced by expert biologists. We measured three different metrics: (1) Dice coefficient quantifies the overlapping between the prediction and the ground-truth, (2) Haussdorf distance indicates the maximum distance between them and (3) the recall (or sensitivity) refers to the fraction of root pixels retrieved over the total amount of root pixels. Quantitative results are included in Table 1. Based on these results, we chose two models, depending on whether we aim at having a faster or more accurate method:

- Fast method: The fastest models are the proposed UNet [6] variants, requiring up to half a second to process a high resolution image using a standard GPU. These models have lower parameter complexity compared to state-of-the-art architectures like SegNet [7] and DeepLab [8], which explains the lower running time. Among the fast UNet models, we observed that the proposed Deeply Supervised ResUNet (DSResUNet) shows a significantly lower value for Hausdorff distance, while keeping equivalently good Dice and Recall. The proposed DSResUNet architecture (depicted in Figure 1) combines residual blocks [9] with deep supervision [10], improving the results of a standard UNet with a minimum increase in model complexity.
- Accurate method: We proposed to combine all the implemented architectures into a single ensemble method, increasing model diversity by creating an ensemble of multiple models and architectures [11]. This ensemble of deep models increased the running time by a factor of 9, but achieved the best performance across all metrics, outperforming state-of-the-art models like SegNet and DeepLab.

**Table 1.** Quantitative evaluation for the different CNN architectures compared in this work. We measured the Dice coefficient, recall and Hausdorff distance for dense root segmentation task. We compared state-of-the-art models (including UNet, ResUNet, SegNet and DeepLab) and compared with the proposed Deeply Supervised ResUNet and the ensemble of multiple models and architectures. On the one side, we found that our Ensemble of Multiple Models and Architectures produced equal or more accurate results than the rest of the models in terms of Dice and Haussdorf, at the expense of increasing the processing time. On the other side, the proposed Deeply Supervised ResUNet is fast (less than half a second), shows a significantly lower value for Hausdorff distance than the other fast models, while keeping equivalently good Dice and Recall.

| Model | Dice |  | Recall |  | Hausdorff Distance (mm) |  | Time | # Params |
| --- | --- | --- | --- | --- | --- | --- | --- | --- |
|  | Thresh | CRF | Thresh | CRF | Thresh | CRF |  |  |
| UNet | 0.769±0.048 | 0.774±0.044 | 0.871±0.044 | 0.830±0.056 | 10.25±7.45 | 9.39±7.94 | 0.29s | 488.212 |
| ResUNet | 0.768±0.050 | 0.770±0.047 | 0.862±0.046 | 0.823±0.057 | 8.83±6.71 | 7.53±5.91 | 0.33s | 505.046 |
| Deeply Supervised ResUNet (our) | 0.769±0.048 | 0.772±0.045 | 0.861±0.044 | 0.815±0.057 | 8.14±7.34 | 6.95±5.42 | 0.49s | 532.336 |
| SegNet | 0.768±0.043 | 0.773±0.040 | 0.862±0.044 | 0.824±0.053 | 7.42±6.40 | 6.81±5.65 | 1.49s | 29.460.450 |
| DeepLab | 0.666±0.055 | 0.609±0.079 | 0.763±0.077 | 0.600±0.113 | 7.58±7.79 | 7.56±7.52 | 1.86s | 58.009.410 |
| Ensamble (our) | 0.772±0.048 | 0.774±0.044 | 0.864±0.044 | 0.804±0.061 | 6.68±5.08 | 6.45±4.98 | 4.5s |  |

ChronoRoot implements both fast and accurate variants, giving the users the opportunity to decide according to their requirements. In this study, we used the fast method based on the proposed Deeply Supervised ResUNet model, which offered a good trade off between running time and accuracy. We apply several post-processing steps after segmentation, which are independent of the CNN model. We first apply a Conditional Random Field (CRF) [12, 13] model to improve the homogeneity of the labels assigned to neighboring pixels. Then, we enhance the temporal consistency of the segmentations by considering its weighted average. These steps serve to remove spurious segmentations by analyzing a temporal sequence of images, which ultimately translates into generating more stable phenotypic measurements. A graph structure is then constructed where every node is assigned a class label indicating whether it is associated with the plant seed, main root, lateral root, bifurcation or root tip. Temporal consistency on the graph structures is finally improved by tracking the labeled nodes and solving conflicting cases. A more detailed description of these steps can be found in the Methods section. After the graph generation process, we proceed to extract phenotypic features for RSA characterization.

### Temporal dimension of traditional and novel RSA parameters

We analyzed temporal sequences of plant roots growing under different conditions. In order to assess the potential of Chrono-Root, we decided to compare RSA of *Arabidopsis thaliana* ecotype Col-0 grown under two distinctive photoperiods, i.e. long day (LD; 16 h of light, 8 h of dark) or continuous light (CL; 24 h of light). Light availability and photosynthesis in the shoot determine the amount of sugar transported to the roots, thus modulating underground plant growth. Moreover, ample evidence suggests that root developmental plasticity depends on the light environment, involving a more sophisticated impact on endogenous signaling pathways [14].

Traditional parameterization of RSA expanded to temporal dynamics revealed the progression of root growth under continuous light (CL) and long day (LD) conditions. A representation of root automatic segmentation is shown in Figure 1 (see Methods and Figure 6 for more details). Our experiments show that main root (MR) length, the sum of lateral root (LR) length and the resulting total root system (TR) begin to differ between conditions at approximately 200-250 h (8-10 days) after germination (Figure 2A, B and C), together with LR number (Figure 2D). Notably, root growth was not only faster under CL, but also resulted in a different RSA, exhibiting a higher density of LRs and a lower component of the MR over the total root system (Figure 2E and F). Notably, between 250 h (10 days) and the end of the experiment (336 h, 14 days), the contrast between both photoperiods increased gradually in every measured parameter, hinting at a temporal reorganization of root development under different light conditions.

**Figure 2.**
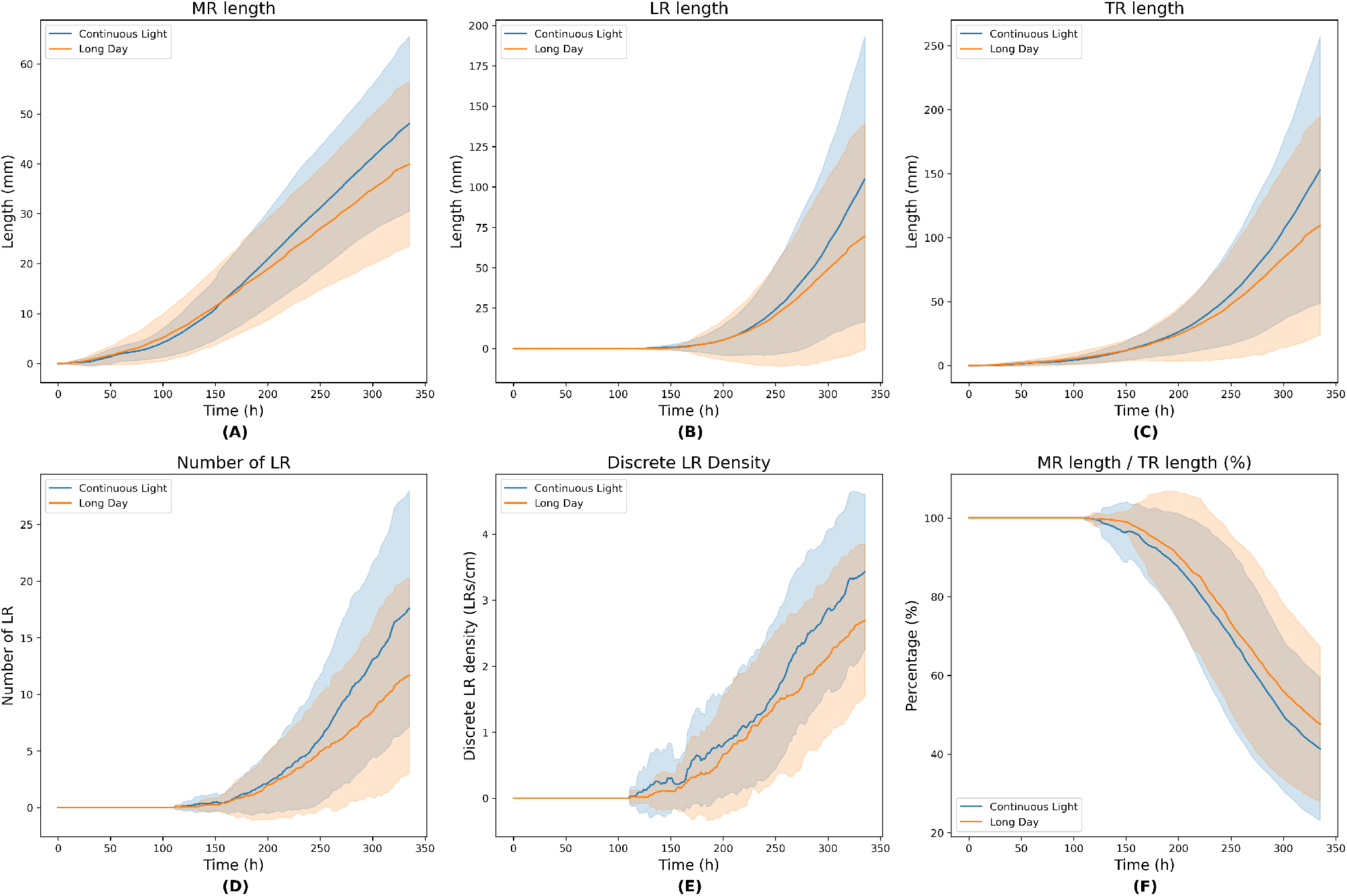
Traditional RSA parameters expanded to the temporal dimension. A. Main root (MR) length; B. The sum of all LRs length; C. Total root (TR) length, representing the sum of LR total length and MR length; D. Number of lateral roots (LRs); E. LR density, expressed as LR number / MR length; and F. MR component of the RSA, expressed as MR length / total root (TR) length, which is the sum of MR and LRs. Data is shown for plants grown under continuous light and long day. The lines indicate the mean, and the shadows represent the standard deviation (SD) throughout the experiment.

Based on the information derived from temporal phenotyping, we explored in more detail the reconfiguration of RSA under alternative photoperiods. We identified the time point at which the sum of LRs length equals the MR length as a novel parameter of RSA dynamics (Figure 3A). However, no significant difference was observed in the distribution of individual time-length points of plants grown under CL or under LD (Figure 3B). The analysis of the relationship between the MR and LRs along time, determined by the difference between both measurements (MR-LRs) aligned to the time point at which MR and LRs are of the same length (time 0) is shown in Figure 3C. It reflects that the difference between MR and LRs tends to have a significantly larger absolute value for plants grown under CL than under LD. Moreover, we extracted different indicators to analyze the dynamics of these curves. Figure 3D shows the approximate derivative (computed by means of finite differences) at the time point at which MR and LRs are of the same lengths (time 0). These differences are not statistically significant according to a Mann Withney U test. However, when extending the analysis to the full +/−24hs range by fitting a linear function to every curve from Figure 3C and plotting the corresponding slopes (Figure 3E), we found strong differences in the distribution (statistically significant according to a Mann Withney U test) for both photoperiods. This novel time-related parameter reflects the dynamics of root growth by determining how long it takes for the system to be composed of more LRs than the MR.

**Figure 3.**
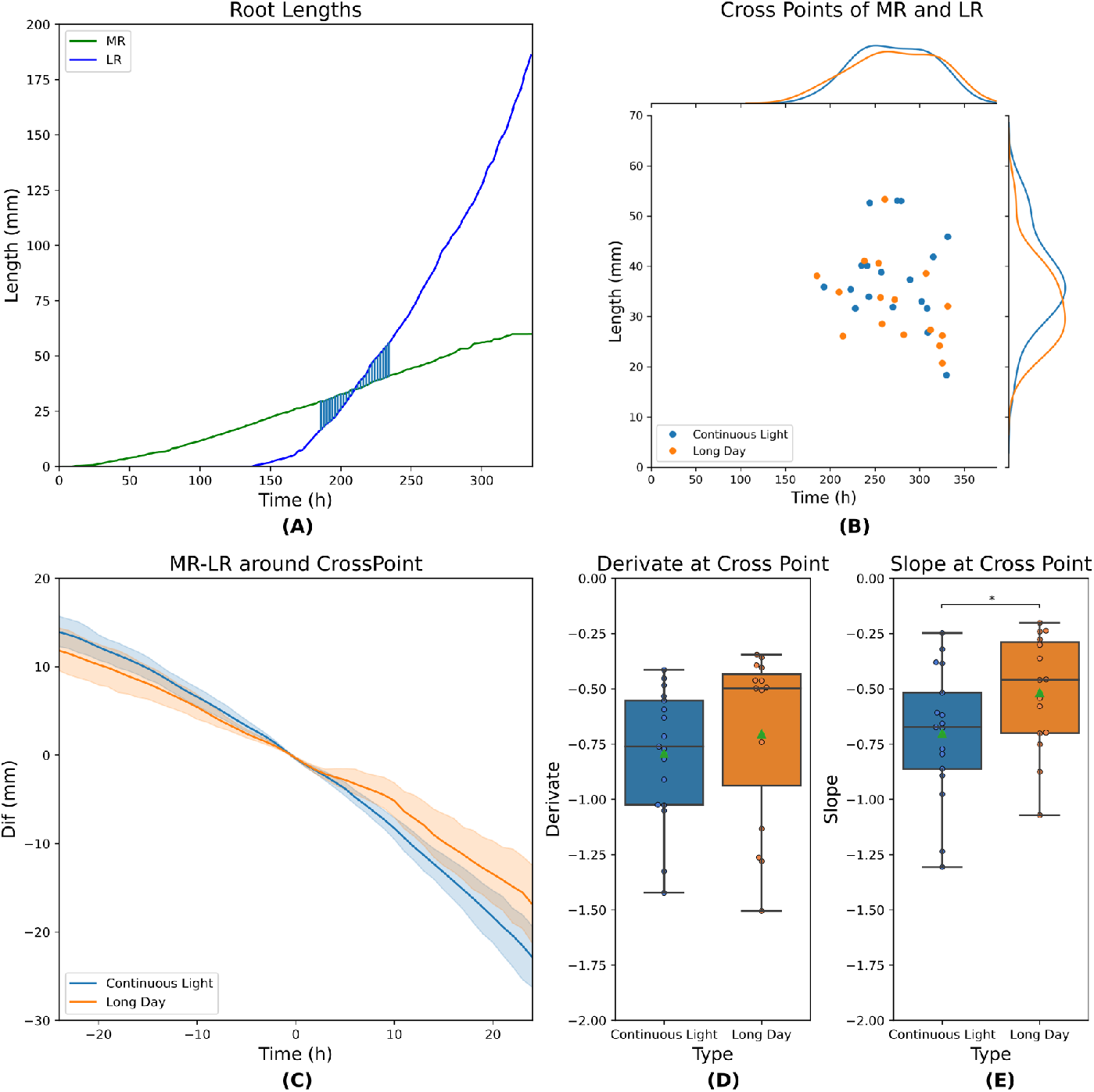
Novel RSA parameters analyzed along time. A. Example of one individual main root (MR) and the sum of lateral root (LR) length along time, revealing the time point of the intersection between the two curves. B. Distribution of the intersection points for all individuals from both conditions (continuous light, CL, and long day, LD). At the top and on the right, the distribution of both populations are represented. For this experiment, no significant difference was observed with respect to the intersection time point. C. The difference between the MR length and total LRs length at each time point was calculated and aligned around the point of MR-LRs = 0 mm for each individual considering +/−24hs. D. Distribution of the derivative value for the individual curves from panel C at time of equal length of MR and total LRs. The difference between the means is not statistically significant according to a non-parametric Mann Whitney U test (p-value > 0.05). E. For every individual, the tendency of the MR-LR curves shown in panel C was determined by fitting a linear function to every curve, considering +/−24hs. The boxplot shows the distribution of the slope of the fitted curves, revealing a clear difference between LC and CL. Difference between the means is statistically significant according to a non-parametric Mann Whitney U test (p-value < 0.05).

In order to assess the impact of RSA reconfiguration on the area explored by roots under distinct photoperiods, we calculated the dynamic convex hull for each subset of plants. Interestingly, the observation of the convex hull resulting from the overlap of all individuals grown in the corresponding conditions reveals an extended high density of LRs along the MR axe at the end of the experiment under CL (14 days after germination, Figure 4A, B and C). Notably, the area of the average convex hulls between 8 and 14 days does not differ between CL and LD conditions (Figure 4D). Nonetheless, the quantification of the sum of LRs length over the convex hull area indicates that the density of LRs is higher under CL between 10 and 14 days (Figure 4E). Collectively, our analyses indicate that global LR length increases under CL as a result of more numerous LRs growing simultaneously, although the area explored by the RSA does not differ between the two photoperiods. Thus, the global density of the resulting RSA is higher under CL.

**Figure 4.**
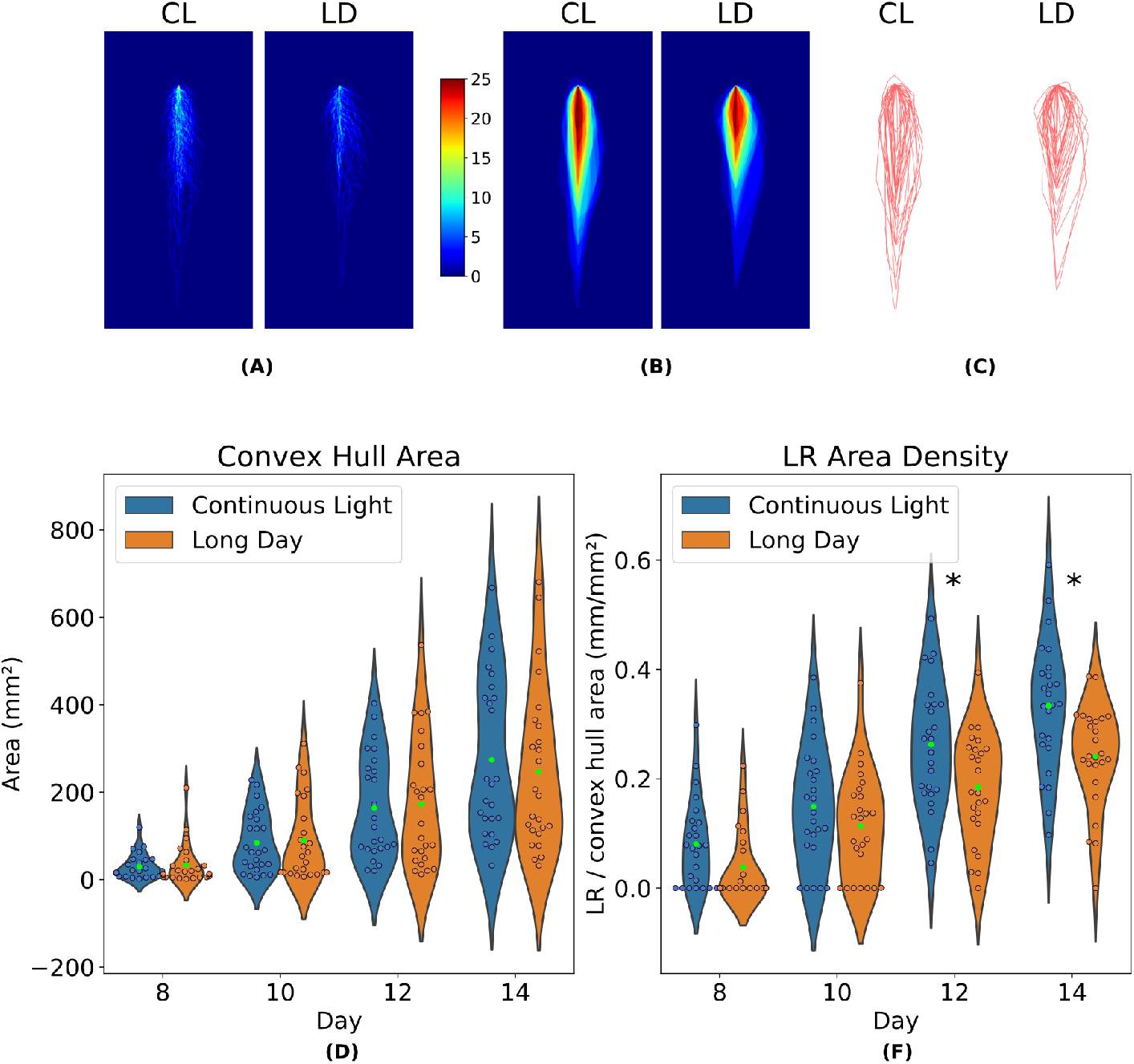
Area and density of the RSA analyzed along time. A. Overlapped segmentations of the whole root system at 14 days after germination. Blue background indicates no roots. The brightness of the signal increases as more roots occupy the same position. B. Same as A, represented as a heat map of the convex hull extracted for each individual. C. Overlapped convex hull contours for each condition. D. Average convex hull area for different time points under CL or LD, represented as violin plots. The mean is indicated as a green point. E. LR density calculated as the sum of LR length over the area included in the respective convex hull. The distribution of each population for the corresponding time points is shown as violin plots. The green points indicate the mean. Asterisks indicate that the difference is statistically significant. We used Shapiro-Wilk test to assess gaussianity, Levene test to confirm equal variances and t-test to confirm that differences between the means of both populations are statistically significant (p-value < 0.01).

### Novel speed-based parameters derived from temporal phenotyping

The information derived from the temporal dimension of traditional and novel RSA parameters indicated that the difference in root growth rate became broader throughout time under CL vs. LD. It has been shown that the *Arabidopsis thaliana* MR exhibits an oscillating growth which likely depends on the lunisolar tide [14, 15] and light-associated carbon partitioning [16]. Therefore, based on the segmentations obtained with our deep learning models, we calculated the growth speed throughout the experiment in both conditions, showcasing how novel speed-based parameters can be derived via ChronoRoot. MR speed grew steadily until approximately 150 h under LD and 200 h under CL post germination, and the average maximum speed reached in CL was higher than in LD (Figure 5A). Strikingly, the difference in the growth speed of the global root system (TR) between the two conditions became increasingly larger since the moment when the speed of the MR was stabilized (Figure 5B), hinting at a different acceleration rate between conditions. The observed root growth dynamics further supports the rising relevance of LRs as a main component of RSA throughout time.

**Figure 5.**
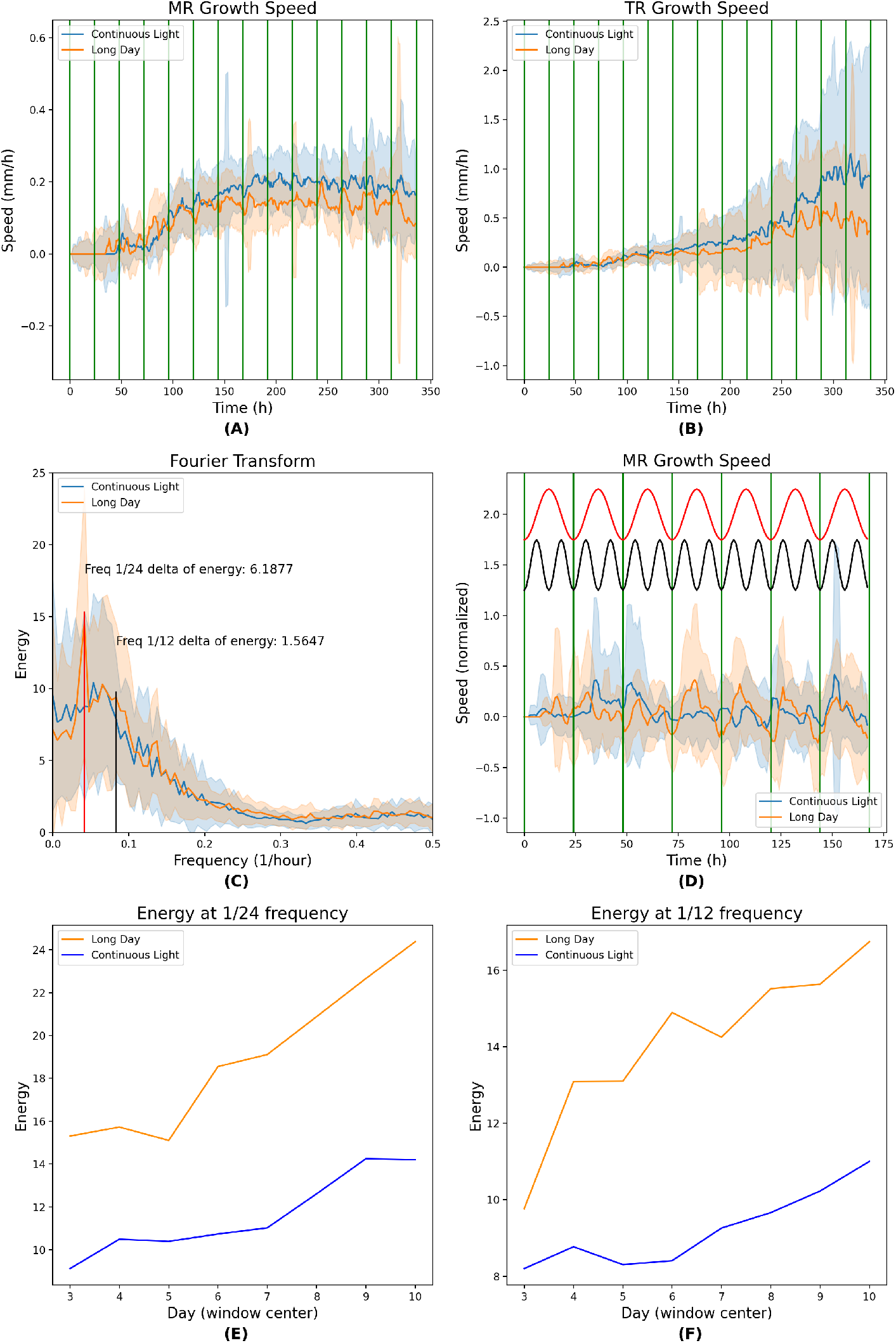
Novel time-derived parameters of RSA. A. Main root (MR) and B. Total root (TR) growth speed along time. C. Fourier Fast Transform of the growth speed signal of MR. The largest energy differences are indicated in the graph. D. The post-processed (high-pass followed by normalization) MR growth speed showing a 7-day-window centered on day 3. The sine curves corresponding to the frequency 1/12 (in black) and 1/24 (in red) found in C are indicated at the top. Note the correlation between the LD growth speed oscillation and the two components 1/12 and 1/24. E. The energy at 1/24 frequency calculated in a 7-day-window centered at consecutive time points for MR. F. The energy at 1/12 frequency calculated in a 7-day-window centered at consecutive time points for MR.

Notably, the analyses of growth speed uncovered an oscillating behavior in both conditions (Figure 5A and B). To better understand the different growth patterns exhibited by LD and CL conditions, we performed a Fourier decomposition of the growth speed signals. Fourier transform decomposes functions depending on time into functions depending on frequency. In other words, the Fourier transform of a given function describes how much of any given frequency is present in the original signal. When comparing growth speed signals, analyzing their Fourier spectrum helps us to see how much this signal correlates with particular oscillation frequencies. For example, if high Fourier coefficients are associated with the frequency 1/24h, it means that the plant growth speed follows a daily oscillation (corresponding to what is known as circadian rhythm). Differences in the Fourier coefficients at a given frequency between growth conditions would indicate an alteration in the oscillatory pattern of plant growth. A Fast Fourier Transformation (FFT) of the signal of MR growth speed in CL vs. LD revealed a major energy difference in the components corresponding to the frequencies of 1/24h and 1/12h, respectively (Figure 5C). Remarkably, these two components distinguish circadian and ultradian rhythms displayed by plants grown in LD, with a pronounced local minimum of the growth speed at 1/24h and a minor local minimum at 1/12h (Figure 5D). Strikingly, the most pronounced differences revealed by FFT (Figure 5C) served to uncover a root growth clock-related disorder suffered under CL, coinciding with a blurred daily oscillation of growth speed, in comparison with the corresponding sine curves (Figure 5D; a detailed comparison of MR, LR and TR growth speed analyses is shown Supplementary Figure 1). Although an oscillating behavior can be observed under CL towards the end of the experiment (Figure 5A and B), the energy at 1/24 and 1/12 frequencies was higher under a LD photoperiod throughout the complete time lapse analyzed (Figure 5E and F). Notably, the difference between conditions of the TR oscillating speed of growth is mainly due to the MR contribution (Supplementary Figure 2). Altogether, our study of wild-type *Arabidopsis thaliana* Col-0 plants growing under alternative photoperiods using ChronoRoot served to reveal novel temporal parameters of root development, notably including clock-related features depending on the light environment.

### 3D-printed device for temporal image acquisition

The ChronoRoot device is an affordable and modular imaging system based on 3D-printed and laser cut pieces and off-the-shelf electronics (Figure 1.1 and 1.2). Each module consists of a Raspberry Pi (v3)-embedded computer controlling four fixed-zoom and fixed-focus cameras (RaspiCam v2), and an array of infrared (IR) LED back-light. In between each camera and the corresponding IR array, there is a vertical 12 × 12 cm plate for seedling growth, allowing automatic image acquisition repeatedly along the experiment without any modification or movement of the imaging setup. The four-plate module is small (62 × 36 × 20 cm) and can be placed in any growth chamber. The different parts of the imaging setup (back-light, plate support and camera) can be positioned along a horizontal double-rail to control the field of view of the camera and accurate lightning. In addition, the camera can be moved vertically. ChronoRoot allows image acquisition at a high temporal resolution (a set of pictures every minute). The use of an IR back-light (850 nm) and optional long pass IR filters (> 830 nm) allow acquiring images of the same quality independently from the light conditions required for the experiment, during day and night.

Each module is connected to the network either by Wi-Fi or Ethernet cable. A web interface allows the control of the device offering live feed of the cameras for field of view and focus setup. The user can program the activation of cameras and IR back-light, starting and ending dates, the time basis for picture acquisition, and finally follow the progression of the experiment. The pictures are saved directly on an external drive plugged on the Raspberry Pi. Once the experimental setup is ready, each module is completely independent from the external environment and the access to the network (for more details see Methods, Supplementary Figures 3-7 and the Supplementary 3D printing and laser cutting files).

## Discussion

The plant phenotype can be defined as the integration of structural, physiological, and performance-related traits of a genotype in a given environment. Plant phenotyping is therefore the act of determining the quantitative or qualitative values of these traits [17]. The advent of novel imaging technologies and image processing have revolutionized plant phenotyping, expanding the frontiers of phenotypic trait measurement. Plant roots have a major role in plant anchorage and resource acquisition while offering environmental benefits such as carbon sequestration and soil erosion mitigation [18]. The growing knowledge linking genetics with functional properties of plant roots is of crucial interest to plant breeding, notably for the design of novel strategies for sustainable agriculture and environmental stewardship in the face of the impending climate change. Whereas high throughput genotyping, sequencing-based genotyping and genomic breeding are behind current agricultural practices in the era of omics technologies, the collection of phenotypic data for a thorough characterization of the RSA is increasingly becoming a limiting factor [19]. Although significant advancements in the application of imaging sensors for high-throughput data collection have allowed comprehensive plant phenotyping [20], the characterization of root traits has been hampered by the inaccessibility of the rhizosphere.

Large and sophisticated phenotyping platforms are deployed worldwide and allow the simultaneous phenotyping of several hundreds of plants (see the International Plant Phenotyping Network[21]). However, their big dimensions and high automatization reserve their implementation on specialized locations and mainly for large phenotyping experiments. Root imaging systems of intermediate complexity, like the one proposed in [22], serve to address the temporal phenotyping of plant roots using vertical plates for plant growth in agarized medium. However, such system still requires a single-axis mobile robot, which implies more expensive electronics and plant chamber space devoted to the equipment setup. In contrast, low-cost ChronoRoot modules can be located easily in already existing facilities without major modifications or permanent movement. The number of modules to be built and used will only depend on the available space and the experimental design (e.g. a few modules for the characterization of given genotypes vs. multiple units for GWAS approaches using tens to hundreds of plant accessions). It also allows to scale the hardware setup according to available funding and experiment requirements progressively. In parallel, the advancement of the Do It Yourself (DIY) movement has promoted the development of a growing number of low-cost phenotyping devices combining 3D-printed, laser cut, captor and micro-controller coming from open-source and open-hardware communities such as Arduino[23] or Raspberry Pi[24]. Successful inexpensive devices have allowed monitoring plant leaves [25], including Phenotiki, an affordable open software and hardware platform for image-based phenotyping of plant aerial organs [26]. More recently, the Phenotiki sensor interface was used to characterize cotton RSA on soil-containing big Rhizoboxes, allowing the determination of basic architectural parameters [27] (like total root area, convex hull area, total root length, etc). In contrast, ChronoRoot allows for a more fine-grained high-throughput temporal phenotyping, e.g. making it possible to distinguish between MR and LRs. Finally, modular rhizotrons of more sophisticated designs (including more expensive cameras and light flashes, precluding observations during the night) also served for RSA characterization of crops [28].

In the last few years, images of the root system from different plant species have been acquired manually using a flat-bed scanner or a camera positioned in front of vertical petri dishes. Root phenotyping is generally performed upon single-time-point images or using several images acquired during growth in time-lapse sequences. More recently, semi-automatic devices and softwares have also helped to increase the efficiency of image acquisition and associated analyses [29, 30, 31, 32, 33, 34]. The great need of throughput in screening experiments to uncover the genetic basis of root development, justifies the use of simplified artificial culture conditions and standardized environments to make the RSA accessible to image acquisition [17]. Software tools use input images of root systems grown under a variety of conditions, including hydroponic and aeroponic systems, agarized medium, paper pouches or soil [33, 32]. Here we propose to use vertical square petri dishes for plant growth on the surface of transparent agarized medium, for automatic acquisition of photographs allowing a high resolution temporal phenotyping of the RSA.

According to Quantitative Plant [35, 36], over 40 image processing softwares are available for root system analysis [37, 33]. RSA parameters are extracted from various types of 2D images captured from agar plates or washed roots extracted from soil. Moreover, 3D RSA reconstruction is possible using X-ray computed tomography [38] or magnetic resonance imaging [39]. Nearly all the reported tools need human input to be operated and retrieve precise numbers. RootTrace [40, 41] for example, which also focuses on high-throughput analyses of root growth, employs traditional image processing and tracking techniques, resulting in a program that can only extract MR length and count the number of emerged LRs. On the contrary, our model relies on deep networks producing a detailed segmentation of the RSA which is then classified into MR and LR, allowing for fine grained measurements like the total length of the LRs, which is not provided by RootTrace. Other tools such as GiaRoots [34] and EZ Rhizo [31] employ simple threshold strategies for root segmentation. In contrast to ChronoRoot, these tools fall short at handling segmentation problems emerging from drops due to water condensation, they require manual human calibration and do not take advantage of the redundancy provided by the temporal resolution of the high-throughput videos to filter out spurious segmentations. Another alternative tool is Win Rhizo [42], a commercial and non-open source tool designed to work with images captured with high resolution desktop optical scanners. Such a requirement makes it virtually impossible to capture high-throughput temporal sequences of growing plants. On the contrary, ChronoRoot is open-source and designed to work with low-cost cameras. Another option is BRAT [43], designed for high-throughput phenotyping of root growth and development. The main disadvantage of BRAT is that it can only handle early root growth, and does not provide measurements for LRs.

The previously discussed methods are mostly based on conventional image processing approaches and extract a limited number of RSA features. Advances in machine learning applied to image analysis allowed partially overcoming these limitations. For example, deep learning techniques have been used to improve the consistency of classic approaches enhancing the quality of root segmentations [44, 45]. Closest to our work is the recent RootNav 2 [46], which is also based on deep learning models and provides fine-grained metrics distinguishing between MR and LRs. However, RootNav 2 does not exploit the redundancy provided by the temporal resolution and follows a different architectural design, which makes ground truth annotations more difficult to obtain and prevents us from training the model with our dataset. Compared to ChronoRoot, RootNav 2 employs a more complex neural network architecture with two output paths: the first one is used to predict root segmentation masks (differentiating between MR and LRs) while the second one produces heat maps associated with root tips. This design choice requires the ground truth annotations to be composed of 3 parts: (1) MR pixel level annotations, (2) LR pixel level annotations and (3) root tip annotations. Conversely, ChronoRoot just requires binary segmentation maps (background vs foreground root) for training, since the MR and LR labeling is performed after segmentation following a depth first search approach on the skeletonized binary segmentation. Thus, our dataset is just composed of images with foreground/background pixel level annotations, which is not enough for training the RootNav 2 model.

## Potential implications

ChronoRoot expands the possibilities for high throughput root phenotyping, which is of major importance for natural variation and GWAS, as well as mutant characterization and screening. Notably, it has been shown that clock-related mutants exhibit a differential oscillating MR growth under alternative conditions [15, 16, 47]. ChronoRoot offers an ideal platform for the identification of genotypes associated with altered clock traits, based on the analysis of spectral features extracted from temporal signals.

Note that the specification of all ChronoRoot hardware components (e.g. camera, etc) is released in this paper. Thus, for anybody installing the system and using the same imaging setup, our software for *Arabidopsis thaliana* analysis should work without further retraining. In case the system fails due to different lighting conditions or hardware components, a minimal fine-tuning of the model by using a few annotated images from the new setup may be required.

## Methods

### Hardware description

An automated imaging setup was designed and built in the shape of an independent module of 62 × 36 × 20 cm (Figure 1). It is aimed at imaging up to four vertical plates either in color or in near-infrared (NIR) lightning. Each module consists of a single board computer (Raspberry Pi) controlling four cameras through a multiplexer module and an array of NIR LED illumination through a relay. The main support of each module is a 620 × 36 × 5 mm acrylic sheet cut using a laser cutter to allow to screw the different parts or let pass strips connecting the camera to the camera multiplexer. Several 3D pieces were designed and printed to place the different components of the module. Each module is separated in four subparts, each of them along a double aluminum axis. This axis allows to adjust the distance of the different parts of the imaging setup: NIR illumination, plate support, camera. The underpart of the module was used to fix the LED AC/DC adaptor, the relay and the computer. The supports under the platform raise and stabilize the module. Supplementary File 1 includes a full description of the components and the steps for the assembly of the device. The 3D-printing and laser cut plans are available online at https://github.com/ThomasBlein/ChronoRootModuleHardware under the CERN Open Hardware Licence Version 2.

### Computational methods

We evaluated different state-of-the-art architectures for image segmentation, and proposed new variants which achieved a good compromise between processing time, model complexity and accuracy, as discussed in the Results section. This segmentation module is followed by several post-processing stages including a CRF post-processing to enhance label homogenity, a temporal consistency refinement step, skeletonization, graph construction and node tracking. ChronoRoot outputs a labeled graph per image indicating which nodes correspond to the seed, main root, lateral roots, bifurcations and the root tips. For each time step, the complete RSA is saved following the RSML format [48].

### Deep learning models for root segmentation

CNNs are representation learning methods with multiple abstraction levels, which compose simple but nonlinear modules transforming representations at one level into a representation at a higher, slightly more abstract level [49]. These models are specially suited for computer vision tasks, in particular for image segmentation [50]. We explored six different convolutional neural network architectures to perform plant root segmentation. Four of them are state-of-the art existing architectures, while the other two were proposed in this work. In what follows, we first present a brief description of the state-of-the-art architectures (namely the UNet [6], ResUNet [51], SegNet and DeepLab [8]) and then discuss the two models proposed in this work.

#### UNet

The first model is a modified lightweight version of the standard UNet [6], which employs a fully convolutional encoder-decoder architecture and produces a dense segmentation map at the pixel level. Based on the original UNet model, we implemented a lightweight version reducing by 4 the number of feature maps per convolutional layer. Skip connections were implemented via summations of the signals in the upsampling part of the network, instead of the concatenation used in the original version. We also replaced the max-pooling layers with avg-pooling, and used an Exponential Linear Unit (ELU) as non-linearity instead of a Rectified Linear Unit (RELU). See Supplementary Table 1 for a detailed description of the implemented architecture.

#### ResUNet

For the second model (ResUNet), we replaced the convolutional layers in the aforementioned UNet architecture by residual blocks [9]. Residual blocks help to prevent the degradation problem which occurs in very deep neural networks by learning residual functions with reference to the layer inputs, instead of learning unreferenced functions. Recent works suggest that residual blocks are effective at segmenting tubular structures like plant roots or roads in a map [51]. See Supplementary Table 2 for a detailed description of the implemented architecture.

#### SegNet

The SegNet architecture [7] is a fully convolutional encoder-decoder neural network, widely adopted by the computer vision community to perform dense image segmentation. The architecture of the encoder is identical to the first 13 layers of VGG-16 [52] and the role of the decoder network is to map the low resolution encoder feature maps to full input resolution feature maps for pixel-wise classification. Differently from the UNet where skip connections are used to propagate the complete feature maps from the encoder to the decoder, the upsampling in the decoder part of the SegNet model uses the memorized max-pooling indices from the corresponding encoder level. Our implementation was based on a publicly available model[53].

#### DeepLab v3

The DeepLab V3 model [8] follows a different approach to generate dense segmentation maps. Differently from the previous models which use skip connections (UNet) or memorized max-pooling indices (SegNet), this model employs atrous convolutions with upsampled filters to extract dense feature maps and capture long range context.

##### Proposed Models

On top of these state-of-the-art architectures, we propose two different CNN models. In the first model, named DSResUNet, we aimed at improving the segmentation accuracy while keeping at the same time a fast lightweight model. In the second case, we focused on increasing the robustness and boosting the accuracy of the segmentation method, at the expense of a more complex model which follows the principle of ensemble learning.

#### DSResUNet

Taking the ResUNet as a baseline model, we propose here a new architecture which combines residual connections and deep supervision [10] to improve the accuracy of the results. Deep supervision integrates additional loss terms which are computed using feature maps from the intermediate CNN layers, instead of the last one only. We concatenated the ResUNet output with the original input image, and processed these feature maps with two additional convolutional layers. This resulted in a cascade of two networks which are trained jointly, where the first one produces an initial segmentation map that is then refined by the second part of the network. We computed two loss terms, one after the output of the standard ResUNet and another one after the additional convolutions. The sum of both terms constitute the loss function used to train the DSResUNet model. See Figure 1 for a graphical illustration of the architecture, and Supplementary Table 2 for a detailed description.

#### Ensemble

Our final segmentation method is an ensemble model. The idea of ensembling is that we can create higher performing models by combining multiple predictors using an aggregation function. One of the most common strategies to implement ensemble models is bagging [54], where the same classifier is trained multiple times using different samples of the training set, and the final output is obtained as the average of the independent predictions. In this work, we followed a different principle which had been successfully applied in the context of medical image segmentation, where instead of combining several instances of the same model trained with different training samples, we combined different models and architectures trained with the same datasets [11]. The idea is to average out the bias infused by individual model configurations, to approximate more reliably the true posterior distribution. In the context of image segmentation, given a dataset *L* = (*x*, *y*)_*i*_ where *x* is an intensity image and *y* the corresponding ground truth segmentation, we aim at learning the underlying conditional distribution *P*(*y*|*x*) which maps input images *x* into segmentation maps *y*. This is commonly approximated by a model *P*(*y*|*x*; θ_*m*_) which has trainable parameters, determined in our case by the neural network architecture. These parameters were learnt so that they minimize a particular loss function (see next section for more details in the loss functions used in our work) using the dataset. Given different architectures (in our case,), we obtained independent estimates of and combined them following [11] approximating the posterior *P*(*y*|*x*) as:

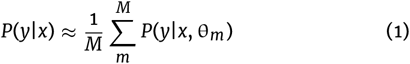

We implemented this ensemble of multiple models and architectures by averaging the predictions of the 5 previous models (UNet, ResUNet, DSResUNet, SegNet and DeepLab v3), obtaining a more robust and accurate segmentation method that significantly outperforms the independent instances.

#### Training details

All the CNN models were trained using binary cross entropy as the loss function, Adam optimizer with default parameters, learning rate of 0.0001 and weight decay = 1e-8 for UNet-like models, 1e-9 for DeepLab and 1e-10 for SegNet. The hyper-parameters were chosen by grid search using the validation data. All models were implemented in Tensor-Flow 1, and the source code is publicly available. The training was done on a standard workstation with Intel(R) Core(TM) i7-8700 CPU, 64 GB RAM and a NVIDIA Titan X graphics processing unit.

Since we are dealing with a relatively small dataset, data augmentation was crucial to achieve good segmentation performance. We implemented online data augmentation through a variety of patch-based augmentation procedures including addition of Gaussian noise, random Gamma corrections to simulate different lighting conditions, artificial blur and horizontal flipping. These transformations were applied to both the images and their corresponding ground-truth segmentation masks. The proposed architectures are all fully convolutional, enabling a patch-based training procedure. As this is a highly unbalanced problem (we have fewer pixels corresponding to root class than background) we implemented the following patch sampling strategy: we sampled patches from random positions centered in root pixels with the same probability as patches centered in background pixels. After performing a grid search of hyperparameters, the size of the training patches was set to 256×256 and we used batches of 8 patches. At test time, we worked with the full resolution images which can be fed to the network and processed by the fully convolutional architectures.

#### CRF Post-processing

The CNN segmentations are postprocessed using a standard fully connected CRF[55]. The CRF operates under the hypothesis that pixels which are contiguous and have similar intensity values should be assigned the same label. We used an efficient publicly available implementation[56] of a dense CRF[13] with Potts compatibility function and hand-tuned parameters θ_α_ = 5 and θ_β_ = 3.

### Graph generation and temporal consistency improvement

The CNN output can be interpreted as a soft segmentation. Since we processed temporal sequences of growing plant roots, we applied a post-processing step to improve temporal consistency using a variation of the weighted trailing average. The current segmentation and an accumulation of the previous ones are averaged to avoid losing parts of the root due to droplets or other type of occlusion. Given the current segmentation *s^t^* at time *t*, and the accumulated mask up to the previous time step *a^t^*^−1^, we compute the current map *a^t^* = *s^t^* + α*a^t^*^−1^. The weight α is chosen depending on the size of the time-step (we used α = 0.9 in our experiments). The aim is to use the root segmentation masks obtained in previous time steps to correct for potentially missing root segments. In our experiments, we processed images every 15 minutes to ensure that the plant has not grown much between two time steps. The average helped to alleviate certain problems caused by root occlusion or water droplets, as the probability maps associated to previous frames act as memory mechanisms, resulting in more stable segmentations (see Figure 6 for a visual example).

**Figure 6.**
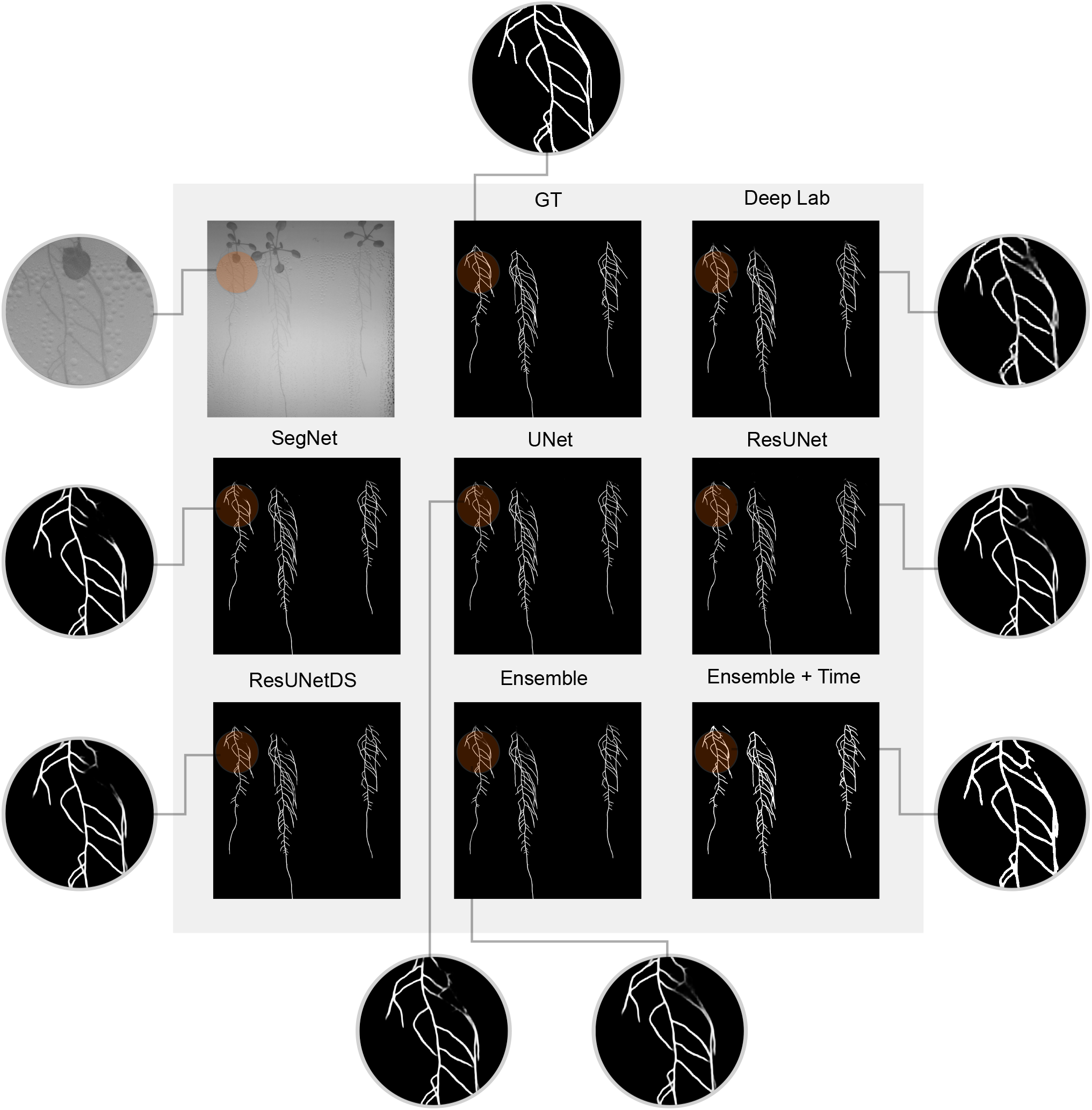
Qualitative segmentation results obtained with the benchmarked methods. We observe how the ensemble and the ensemble with temporal consistency improve the quality of the results, specially in areas with root occlusion.

At this point, as the user selects a Region of Interest (ROI) for each plant, the algorithm starts working one by one. We proceed to threshold the accumulated probability map for the selected plant, perform closing and opening morphological operations[57] to eliminate spurious pixels and then we select the biggest connected component as the root segmentation. Finally, we proceed to skeletonize[58] the segmentation and construct a graph that represents the root system architecture.

We run a depth first search (DFS) algorithm[59] in order to label the bifurcation and end nodes of the unlabeled root graph given by the skeletonized binary segmentation. We use the DFS algorithm, starting from a seed that can be automatically chosen as the top pixel in the plant ROI or manually specified. For assigning labels to the MR, we work based on the assumption that in early growing stages, there will only be a MR with seed (top pixel) and tip (bottom pixel). We then use nearest neighbours for matching the node graphs in the succeeding iterations. As more nodes appear deviating from the MR, they will be added as bifurcation (more than one neighbour) or lateral root tip (one neighbour, different from the MR tip). In case that one LR collides with the main root or another LR, the tip will still be a tip because of the matching process. Following this procedure, labels are assigned for the seed, main root tip, bifurcation and lateral root tip nodes. Node graph matching based on a nearest neighbor criterion was performed between the labeled nodes of successive graphs in the temporal sequence to track the evolution of the root. These graph structures allowed us to extract phenotyping features such as main root length, total lateral roots length or number of lateral roots at every temporal step. By processing the complete temporal sequence for a given root, we can obtain temporal features such as growing speed or information about the root behavior on day-night cycles, enabling the emergence of novel temporal plant phenotypes, as those shown in the Results section. Figure 7 includes several examples of RSAs extracted from images with different levels of complexity. Note that we visualize the graphs using a simplified version where only nodes corresponding to seed, bifurcation and tips are shown and connected. However, it is important to highlight that MR and LR length are computed considering the real length along the labeled skeleton, which are stored as an edge attribute in the simplified graph for optimization reasons.

**Figure 7.**
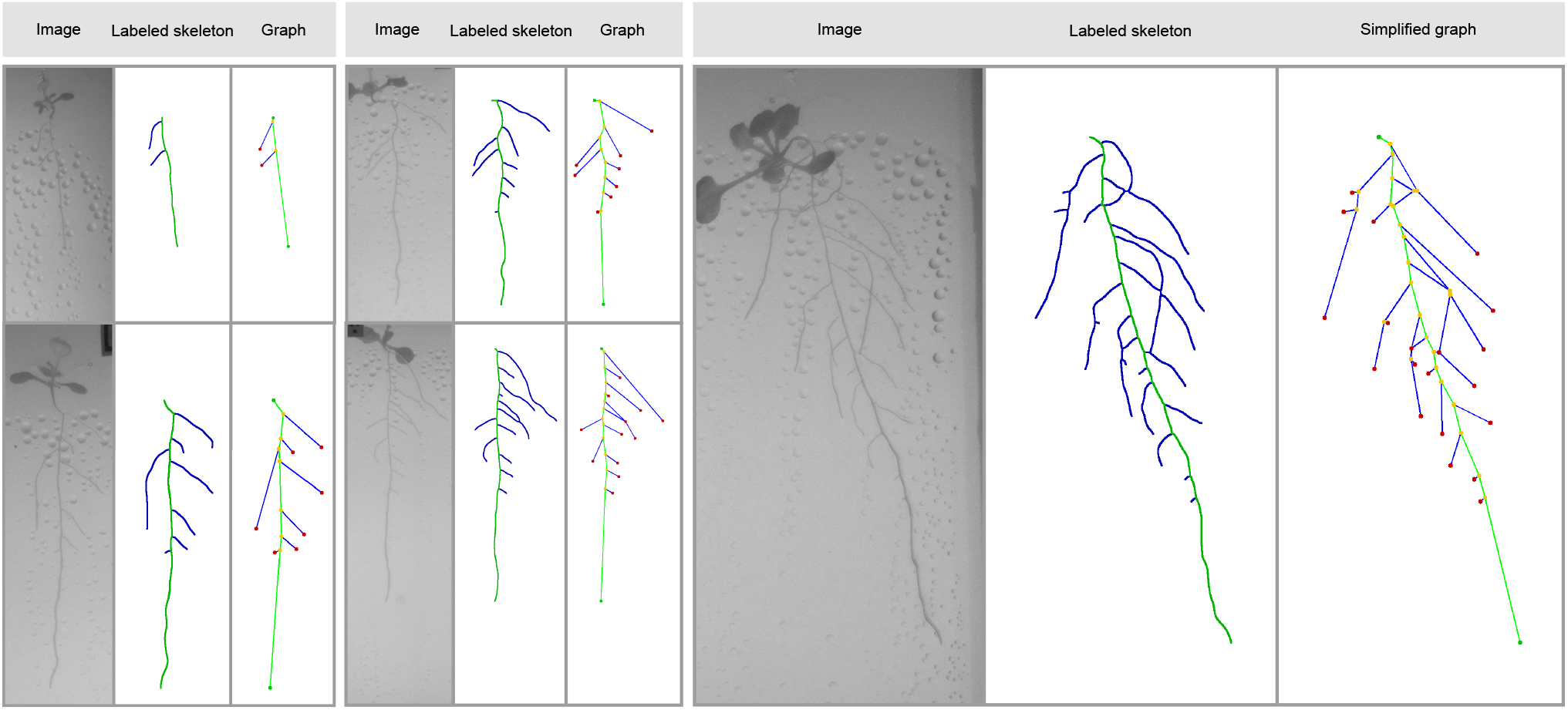
Examples of images, labeled skeleton and simplified graphs corresponding to RSAs exhibiting different levels of complexity. Note that we visualize the graphs using a simplified version where only nodes corresponding to seed, bifurcation and tips are shown and connected. However, since the full skeleton is labeled, the MR and LR length are computed considering the real length along the skeleton.

## Availability of source code and requirements

The source code corresponding to ChronoRoot, namely the deep learning model and the graph generation procedures:

- Project name: ChronoRoot: High-throughput phenotyping by deep learning reveals novel temporal parameters of plant root system architecture
- Project home page: https://github.com/ngaggion/ChronoRoot
- Research Resource Identifier (RRID): SCR_021259
- Operating system(s): Platform independent
- Programming language: Python
- Other requirements: Python > 3.3, Anaconda, TensorFlow 1.15, PyDenseCRF
- License: GNU GPL

The source code corresponding to ChronoRoot imaging controller, namely the web interface to check and set up the image acquisition parameters:

- Project name: ChronoRoot: Module Controller
- Project home page: https://github.com/ThomasBlein/ChronoRootControl
- Operating system(s): GNU/Linux
- Programming language: Python
- Other requirements: NGINX, uWSGI, Python >= 3.5, Flask >= 1.1.0, APScheduler, RPi.GPIO, picamera, WTForms, smbus2
- License: OSI-approved CeCILL-2.1 license

## Availability of supporting data and materials

All data gathered and reported in this study are available as supplementary material. The two datasets of images and annotations described in the “Datasets” section, as well as the 3D printing and laser cutting files are publicly available at https://github.com/ThomasBlein/ChronoRootModuleHardware under the CERN Open Hardware Licence Version 2 - Strongly Reciprocal licence. Supplementary figures and tables referenced in this work, as well as a detailed description of the hardware system are available in the annex Supplementary File 1. Snapshots of our code and other data further supporting this work are openly available in the GigaScience Repository, GigaDB[60].

## Additional Files

**Supplementary File 1** includes the following figures and tables:

- Supplementary Table 1: Detailed description of the UNet architecture implemented in this work.
- Supplementary Table 2: Detailed description of the Residual UNet implemented in this work and the proposed Deeply Supervised Residual UNet.
- Supplementary Figure 1: Novel time-derived parameters of RSA. Supplementary Figure 2: Novel time-derived parameters of RSA.
- Supplementary Figure 3: Low-cost device for automatic image acquisition of plant plates.
- Supplementary Figure 4: LED near-infrared panel front view and back view.
- Supplementary Figure 5: Plate support.
- Supplementary Figure 6: The camera setup.
- Supplementary Figure 7: Electronic connection of a module.

**Supplementary File 2** includes a Video Abstract of this work.

## Declarations

## List of abbreviations

CL: continuous light
CNN: convolutional neural network
CRF: conditional random field
DFS: depth first search
DSResUNet: Deeply Supervised ResUNet
ELU: exponential linear unit
FCN: fully convolutional network
FFT: fast Fourier Transform
GPU: graphical processing unit
GWAS: genome-wide association studies
GT: ground truth
IR: infra-red
LD: long day
LR: lateral root
MR: main root
NIR: near infra-red
RELU: rectified linear unit
ROI: region of interest
RSA: root system architecture
RSML: Root System Markup Language
SD: standard deviation
TR: total root

## Consent for publication

Not applicable

## Author’s Contributions

TB, MC, EF and FA conceived the project. NG and EF designed the deep learning models. NG implemented the deep learning models, ran the numerical experiments and generated the figures. TB, VD, EL and SL designed and built the hardware system. TB and VD implemented the web control interface. TR prepared the plates with *Arabidopsis thaliana* seeds and launched the experiments for image acquisition. AC and NG annotated the images used to train the deep learning models. NG, EF, FA, DM and TB analyzed and interpreted the results. EF, FA, TR, DM, MC, TB, NG wrote the paper.

## Acknowledgements

We would like to thank Fablab Digiscope | LRI | UPSACLAY, and in particular Romain Di Vozzo, for fruitful discussions, his advice in the design and for the access to their digital fabrication equipment. We thank Jean-Paul Bares and Maël Jeuffrard from IPS2 for support and assembling of the ChronoRoot modules. We gratefully acknowledge the support of NVIDIA Corporation with the donation of the Titan Xp used for this research.

## Authors’ information

FA, DM and EF are researchers of CONICET; NG and AC are fellows of the same institution. TB and MC are researchers and VD is an engineer of CNRS. EL and SL are technicians and TR is a fellow of University Paris-Saclay.

