## Supplementary tables and figures for "ChronoRoot: High-throughput phenotyping by deep segmentation networks reveals novel temporal parameters of plant root system architecture"

### Supplementary Files

#### CNN Architectures

| U-Net | | | |
|---|---|---|---|
| **Block** | **Input Dimensions** | **Feature Maps** | **Output Dimensions** |
| Conv / Batch Norm /Elu<br>Conv / Batch Norm /Elu | 256x256 | 16 | 256x256x16 |
| Average Pooling 2x2 | 256x256x16 | | 128x128x16 |
| Conv / Batch Norm /Elu<br>Conv / Batch Norm /Elu | 128x128x16 | 32 | 128x128x32 |
| Average Pooling 2x2 | 128x128x32 | | 64x64x32 |
| Conv / Batch Norm /Elu<br>Conv / Batch Norm /Elu | 64x64x32 | 64 | 64x64x64 |
| Average Pooling 2x2 | 64x64x32 | | 32x32x64 |
| Conv / Batch Norm /Elu<br>Conv / Batch Norm /Elu | 32x32x64 | 128 | 32x32x128 |
| Dropout Layer | | | |
| Upconv | 32x32x128 | 64 | 64x64x64 |
| Elu(Skip Connection (ADD)) | 64x64x64 | | 64x64x64 |
| Conv / Batch Norm /Elu<br>Conv / Batch Norm /Elu | 64x64x64 | 64 | 64x64x64 |
| Dropout Layer | | | |
| Upconv | 64x64x64 | 32 | 128x128x32 |
| Elu(Skip Connection (ADD)) | 128x128x32 | | 128x128x32 |
| Conv / Batch Norm /Elu<br>Conv / Batch Norm /Elu | 128x128x32 | 32 | 128x128x32 |
| Dropout Layer | | | |
| Upconv | 128x128x32 | 16 | 256x256x16 |
| Elu(Skip Connection (ADD)) | 256x256x16 | | 256x256x16 |
| Conv / Batch Norm /Elu<br>Conv / Batch Norm /Elu | 256x256x16 | 16 | 256x256x16 |
| Conv | 256x256x16 | | 256x256x2 |

**Supplementary Table 1.** Detailed description of the UNet architecture implemented in this work.

| DS Residual U-Net | | | | |
|---|---|---|---|---|
| **Block** | **Input Dimensions** | **Feature Maps** | **Output Dimensions** | **Model** |
| Residual Block | 256x256 | 16 | 256x256x16 | **Residual U-Net** |
| Max Pooling 2x2 | 256x256x16 | | 128x128x16 | |
| Residual Block | 128x128x16 | 32 | 128x128x32 | |
| Max Pooling 2x2 | 128x128x32 | | 64x64x32 | |
| Residual Block | 64x64x32 | 64 | 64x64x64 | |
| Max Pooling 2x2 | 64x64x32 | | 32x32x64 | |
| Residual Block | 32x32x64 | 128 | 32x32x128 | |
| Dropout Layer | | | | |
| Upconv | 32x32x128 | 64 | 64x64x64 | |
| Elu(Skip Connection (ADD)) | 64x64x64 | | 64x64x64 | |
| Residual Block | 64x64x64 | 64 | 64x64x64 | |
| Dropout Layer | | | | |
| Upconv | 64x64x64 | 32 | 128x128x32 | |
| Elu(Skip Connection (ADD)) | 128x128x32 | | 128x128x32 | |
| Residual Block | 128x128x32 | 32 | 128x128x32 | |
| Dropout Layer | | | | |
| Upconv | 128x128x32 | 16 | 256x256x16 | |
| Elu(Skip Connection (ADD)) | 256x256x16 | | 256x256x16 | |
| Residual Block | 256x256x16 | 16 | 256x256x16 | |
| Conv (ResUNet OUT) | 256x256x16 | | 256x256x2 | |
| Concat (ResUnet OUT) with original input | 256x256x2 | | 256x256x3 | **Deeply Supervised Residual U-Net** |
| Residual Block | 256x256x3 | 16 | 256x256x16 | |
| Max Pooling 2x2 | 256x256x16 | | 128x128x16 | |
| Residual Block | 128x128x16 | 32 | 128x128x32 | |
| Dropout Layer | | | | |
| Upconv | 128x128x32 | 16 | 256x256x16 | |
| Elu(Skip Connection (ADD)) | 256x256x16 | | 256x256x16 | |
| Residual Block | 256x256x16 | 16 | 256x256x16 | |
| Conv (DSResUNet OUT) | 256x256x16 | | 256x256x2 | |

**Supplementary Table 2.** Detailed description of the Residual U-Net implemented in this work and the proposed Deeply Supervised Residual U-Net.

### Supplementary Figures

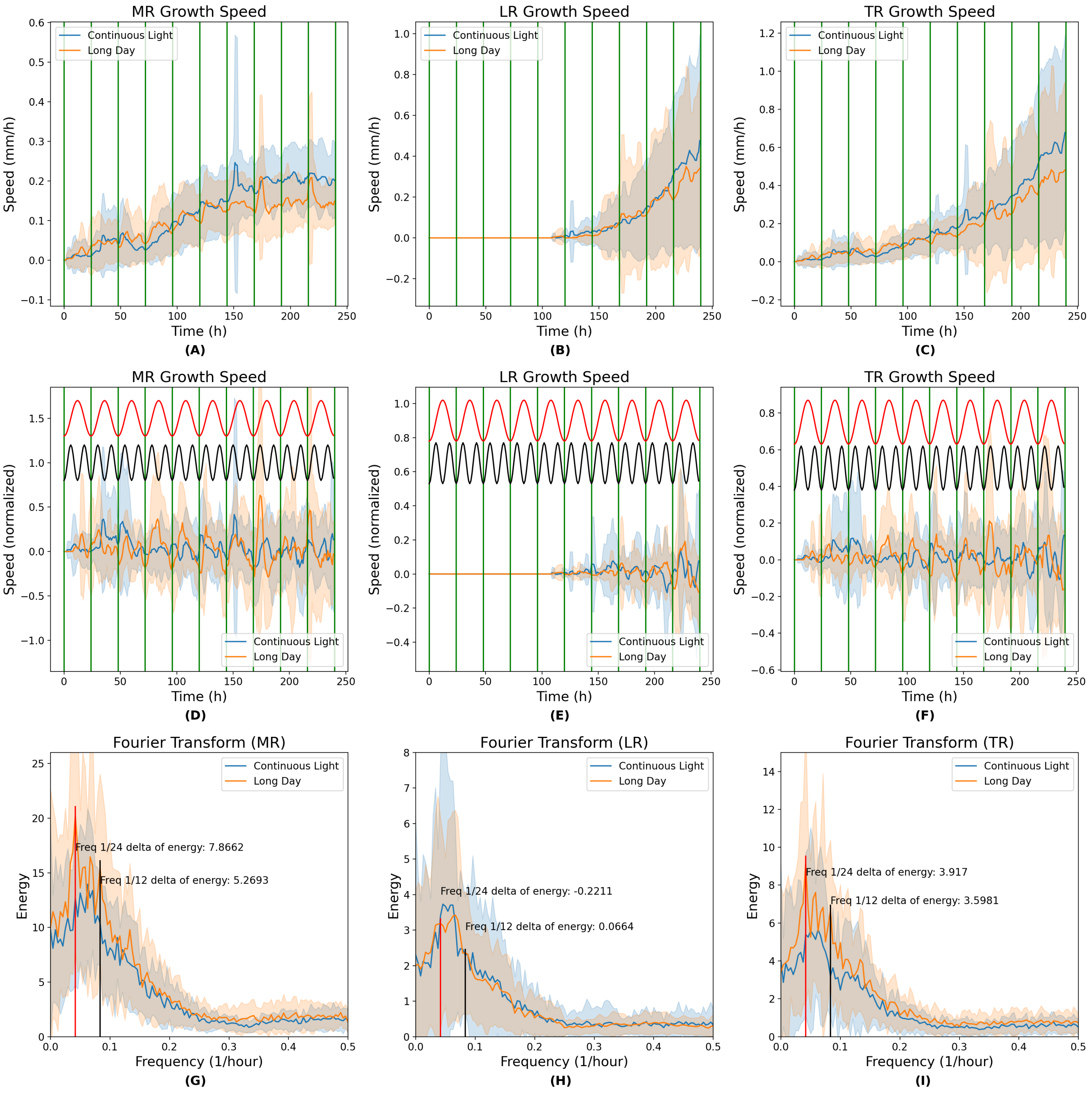


**Supplementary Figure 1. Novel time-derived parameters of RSA.** A. Main root (MR), B. Lateral Roots (LR) and C. Total root (TR) growth speed along time. D-F. Post-processed speed along time (application of a high-pass filter followed by normalization) of (D) MR, (E) LR and (F) TR, respectively. The sine curves corresponding to the frequency 1/12 (in black) and 1/24 (in red) found in D-F are indicated at the top of each panel. G-I. Fourier Fast Transform of the growth speed signal of (G) MR, (H) LR and (I) TR, respectively.

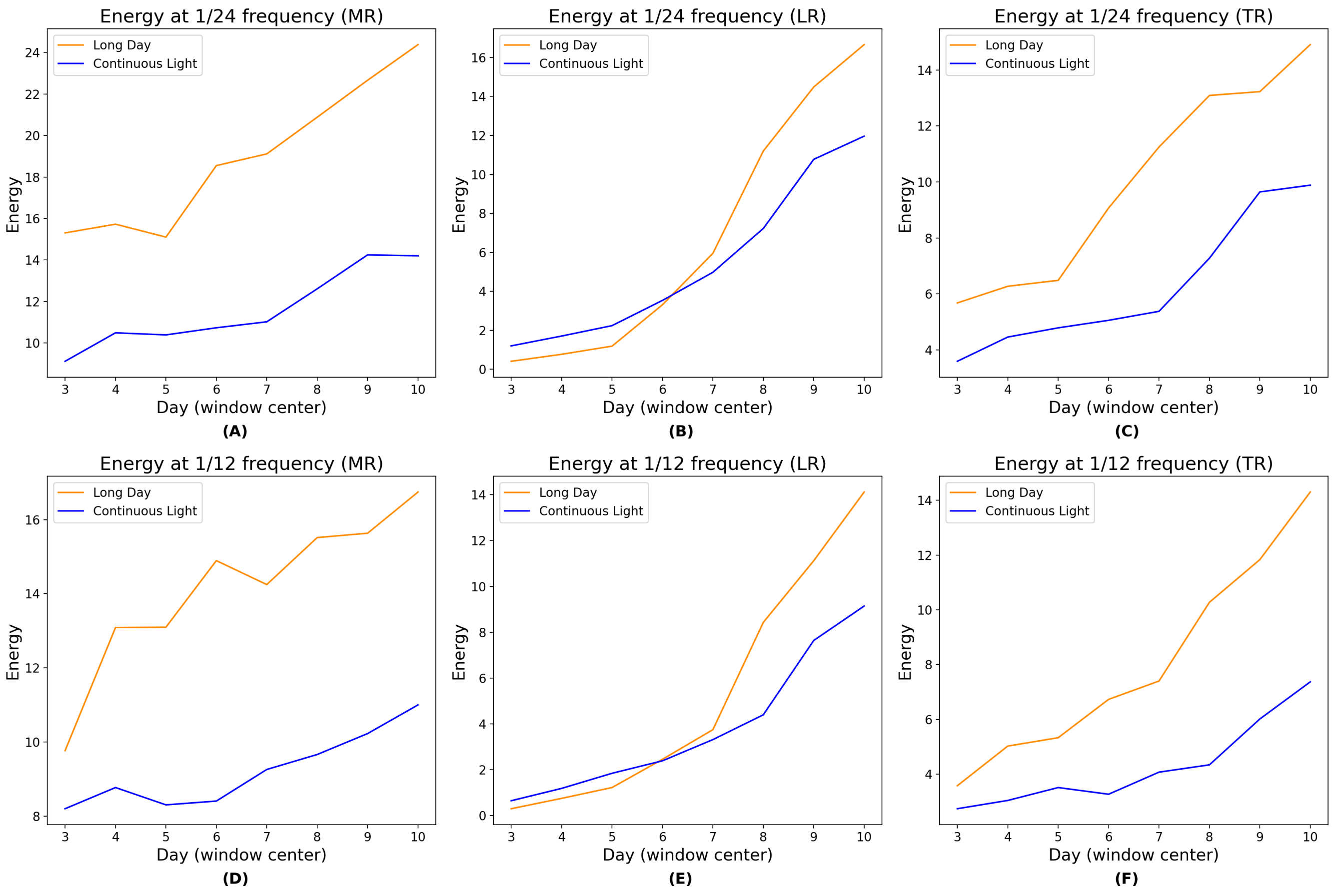


**Supplementary Figure 2. Novel time-derived parameters of RSA.** A-C. The energy at 1/24 frequency calculated in a 7-day-window centered at consecutive time points for (A) MR, (B) LR and (C) TR. D-F. The energy at 1/12 frequency calculated in a 7-day-window centered at consecutive time points for (D) MR, (E) LR and (F) TR.

#### Hardware Description

In this section we include supplementary notes with additional details about the hardware specification. These notes are provided as open-hardware specifications, to encourage other scientists to 3D print and mount the device in their own laboratories.

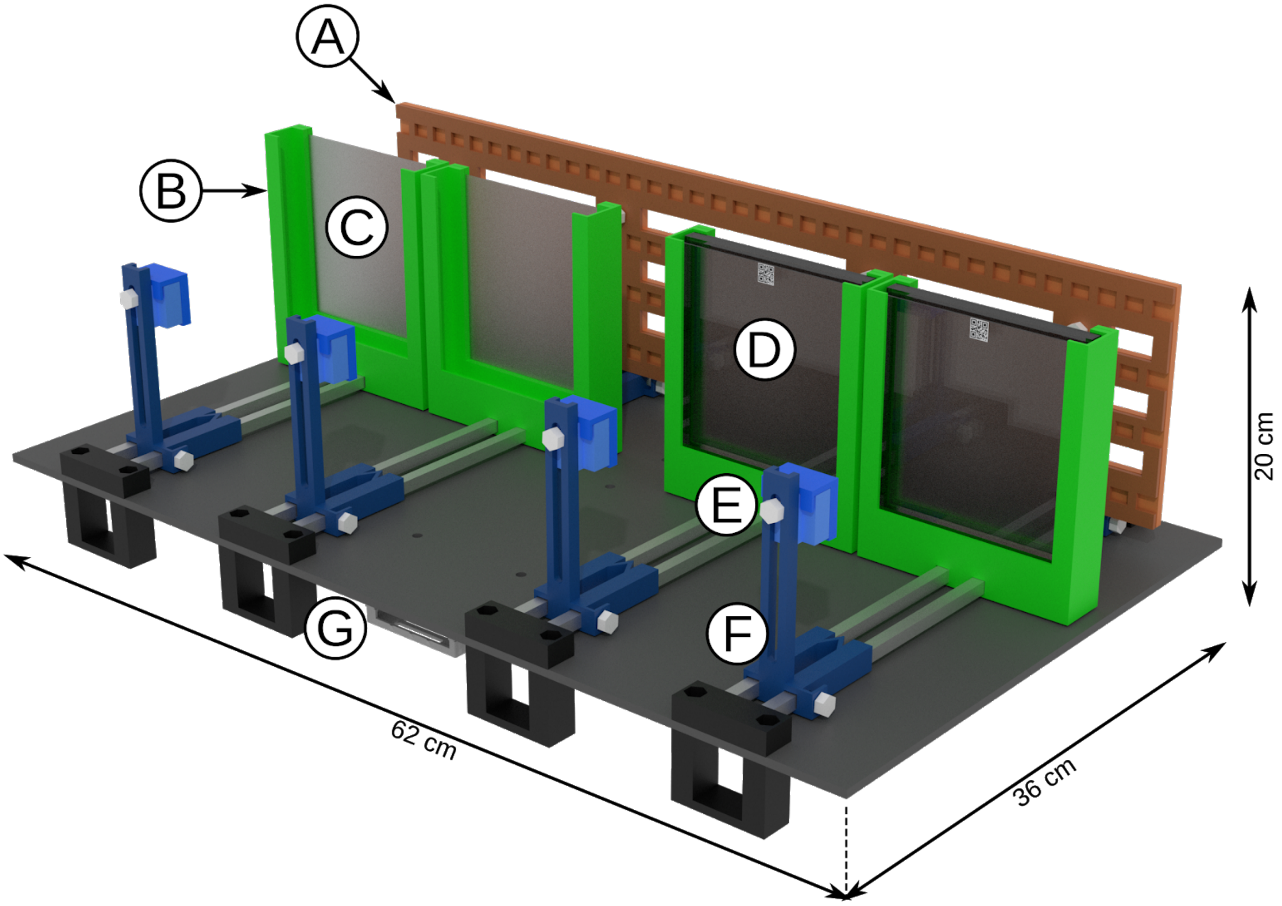


**Supplementary Figure 3. Low-cost device for automatic image acquisition of plant plates.** 3D rendering of a ChronoRoot module. (A) mobile and controlled infrared (IR) backlight (850 nm), (B) mobile plate support, (C) diffusing filter, (D) 12 cm x 12 cm square plate with QR code on the top, (E) Camera case equipped with an IR long pass filter (> 830 nm), (F) mobile camera support, (G) Raspberry Pi computer controlling the module (IR backlight and camera).

The NIR illumination was build with four rows of LED flexible tape (tri-chip SMD5050-150-IR 850 nm, Huake LTD, China) fixed in a sandwich between two acrylic plates (580 mm x 140 mm x 3 mm) laser cuts to allow air flow and prevent the LED strip from ungluing. The four strips of LED were connected in parallel to a 12V AC/DC adaptor fixed under the module. The power supply cable of the adaptor is under the control of relay (Single Relay Board #27115, Parallax Inc), in a box fixed near the adaptor. It allows the control of the NIR illumination by the computer. The LED array is maintained vertically by L-shaped supports able to move along the aluminum horizontal axes as four individual panels in the module, corresponding to the respective plant plates.

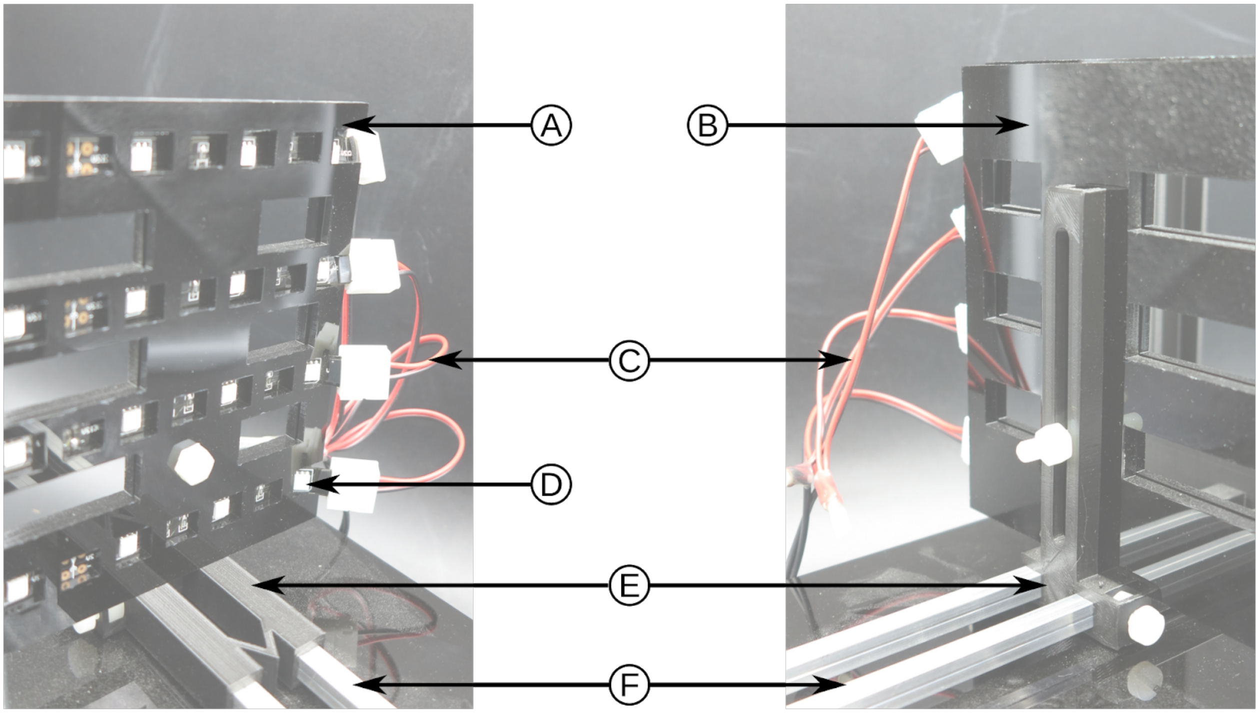


**Supplementary Figure 4. LED near-infrared** panel front view (left) and back view (right). The LED strips (D) are squeezed in between two acrylic panels (A and B) to prevent them from unsticking from the back panel. The strips are linked in parallel to the AC/DC adaptor through strip connector (C). The full LED panel is kept in vertical position via an L-shaped support (E) able to move along the double aluminum axis (F). Acrylic screws and nuts are used to maintain the LED panel on the L-shape support and fixed the horizontal position on the horizontal axis. U-shaped plate holders were designed to hold 12.5 cm x 12.5 cm square petri dishes used to grow the plants. The plate holder carries a diffusion filter (Cinegel R3000 Tough Rolux, Rosco Laboratories Inc.) at the back of the plate to allow homogeneous backlight illumination of the plates by the NIR LED array. The four plate holders can be moved independently along their respective horizontal axis.

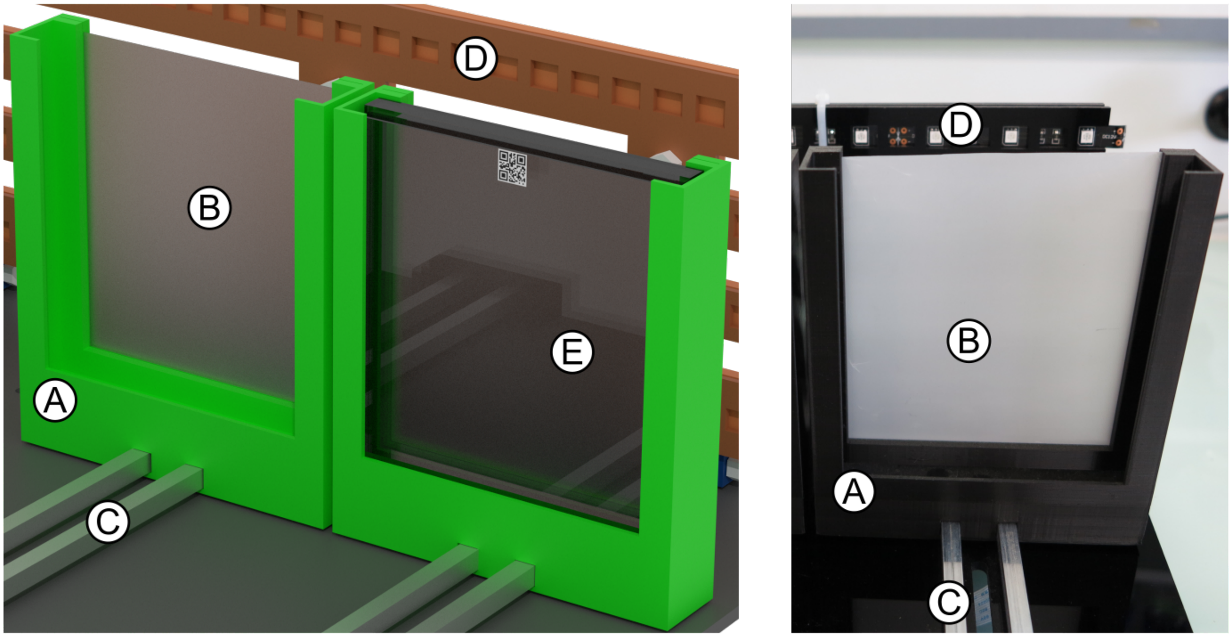


**Supplementary Figure 5. Plate support.** Annotated 3D rendering (left) and picture (right) of the plate support. The U-shaped plate support (A) carries a diffusion filter at the back (B) allowing an homogeneous near-infrared illumination. Each support moves horizontally on the double axis (C). A classical 12.5 cm x 12.5 cm square plate fit into the plate support (E). Each camera is a regular Raspberry Pi NoIR V2 module (Sony IMX219 8-megapixel sensor) which is able to capture NIR wavelengths in addition to the classical visible spectrum. It is positioned in a box that can be moved vertically on an L-shaped support which can be shifted along the horizontal axis. The focus of the camera is adjusted manually before the start of the experiment by turning the objective lens using the provided crown. To be able to record pictures exclusively from the NIR spectrum, an IR long pass filter (12.5 mm diameter, RG830 Schott AG) can be positioned in front of the camera to exclude light below 830 nm and provide consistent images independently of the lightning of the growth chamber.

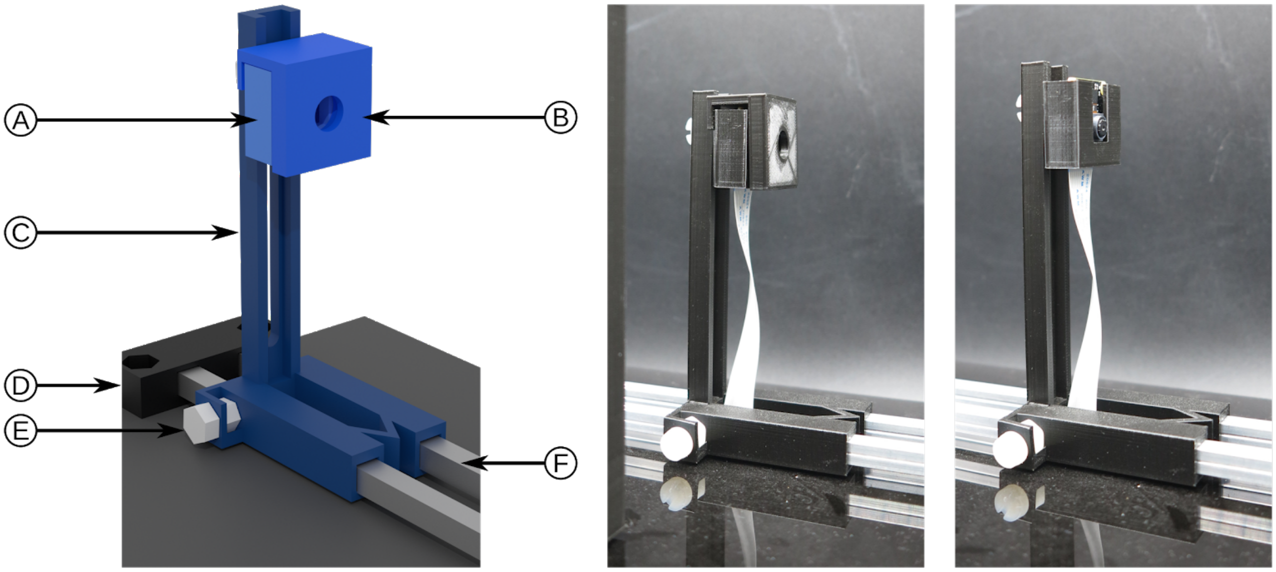


**Supplementary Figure 6. The camera setup.** (Left) annotated 3D rendering; (middle) camera on a module carrying a filter; (right) camera on a module without the filter. A Raspberry Pi NoIR V2 module is enclosed in a 3D printed box (A) that could carry an optional filter (B). The box can be moved vertically on an L-shaped support (C) which itself is able to move along a double aluminum axis (F); The position of the L-shape support and of the camera box are secured by acrylic screw and nut (E for L-shaped support). The double aluminum axis is attached to the main board by a 3D printed part (D). The computer is a Raspberry Pi 3 model B with an additional camera multiplexer module (IVPort V2 Raspberry Pi Camera Module V2 Multiplexer, Ivmech Mekatronik & İnovasyon Ltd.) and allows to connect four cameras. It is boxed in a cage fixed under the main board. The computer runs the Raspbian GNU/Linux operating system and a python web application was developed to control the different components of the module. The camera multiplexer module is controlled through the ivport_v2 library (https://github.com/ivmech/ivport-v2), the control of relay of the NIR backlight through pins 30 (ground) and 32 (GPIO 12) of the Raspberry Pi GPIO through the RPi.GPIO library (https://pypi.org/project/RPi.GPIO/). The web interface was developed using the Flask web framework (https://flask.palletsprojects.com), uWSGI (https://uwsgi-docs.readthedocs.io/), Twitter Bootstrap for the frontend (https://getbootstrap.com/). The scheduling of the task was managed using the APScheduler library (https://apscheduler.readthedocs.io). The source code of the application is available at https://laforge.ips2.u-psud.fr/thomas.blein/kinematicrobot.

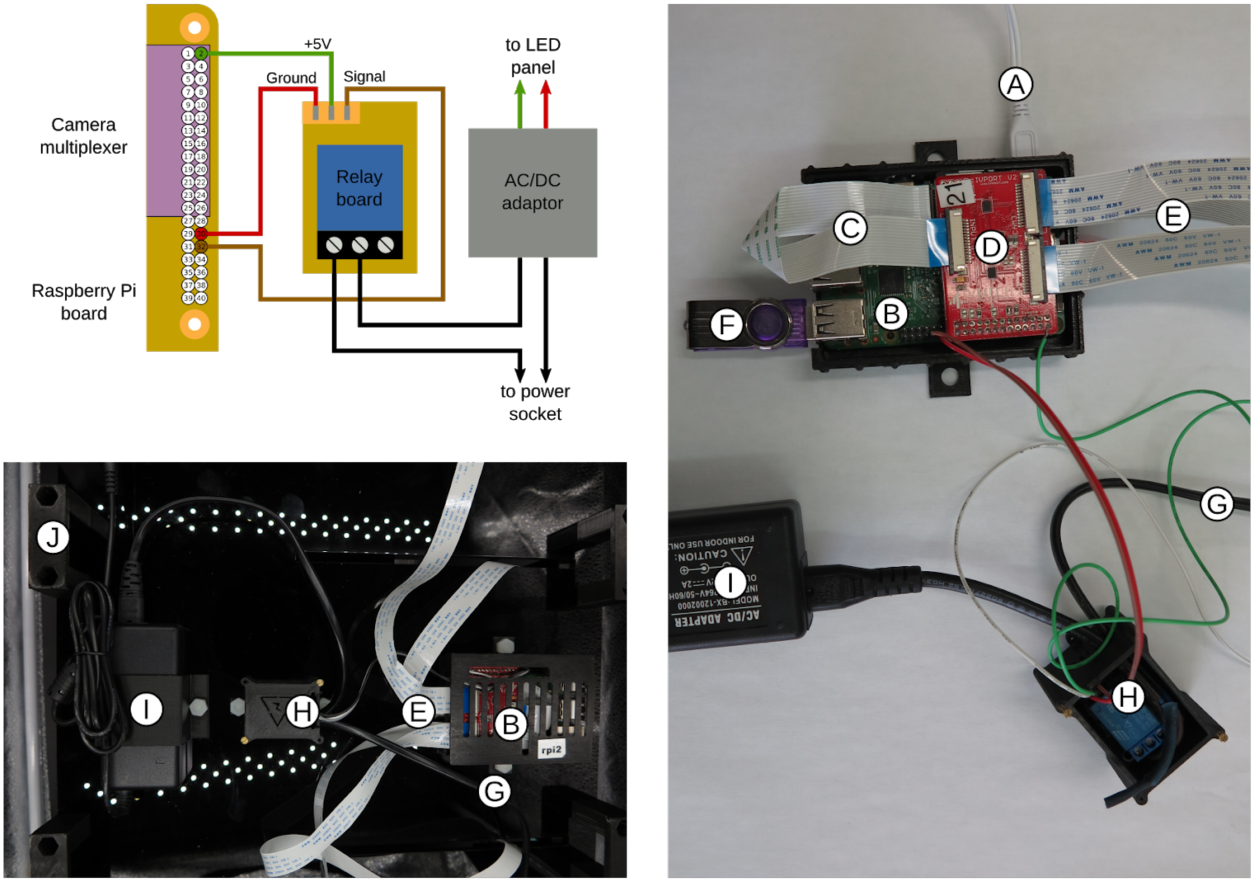


**Supplementary Figure 7. Electronic connection of a module.** Schematic electric connection between the Raspberry Pi GPIO, the camera multiplexer, the relay board and AC/DC adaptor (up left). Real view of the connection with different part boxes open (right). The Raspberry Pi (B) is connected to a USB power adaptor (A). The camera multiplexer module (D) is plugged on the Raspberry Pi GPIO port and connected to the Raspberry Pi camera by a strip (C). Each camera is connected to the multiplexer by a strip (E). The pictures are saved on a USB stick (F) or a USB hard drive directly connected to the Raspberry Pi. A relay board (H) is connected to the Raspberry Pi GPIO and controls the main power (G) of the AC/Dc adaptor powering the LED strip. The final organization of the controlling part under the module main board raised by foots (J).
